# High-fidelity continuum modeling predicts avian voiced sound production

**DOI:** 10.1101/790857

**Authors:** W. Jiang, J.H. Rasmussen, Q. Xue, M. Ding, X. Zheng, C.P.H. Elemans

## Abstract

Voiced sound production is the primary form of acoustic communication in terrestrial vertebrates, particularly birds and mammals, including humans. Developing a causal physics-based model that links descending vocal motor control to tissue vibration and sound requires embodied approaches that include realistic representations of voice physiology. Here we first implement and then experimentally test a high-fidelity three-dimensional continuum model for voiced sound production in birds. Driven by individual-based physiologically quantifiable inputs, combined with non-invasive inverse methods for tissue material parameterization, our model accurately predicts observed key vibratory and acoustic performance traits. These results demonstrate that realistic models lead to accurate predictions and support the continuum model approach as a critical tool towards a causal model of motor control of voiced sound production.

## Main text

During voiced sound production laryngeal vocal folds (VFs) or analogous syringeal structures exhibit self-sustained vibrations through fluid-tissue interactions and elastic restoring forces. VF vibrations mechanically convert expiratory airflow into pulsatile airflow, which causes air pressure oscillations constituting the acoustic excitation of the system (*1*–*4*). This physical framework is known as the myoelastic-aerodynamic theory (MEAD) (*2*, *3*) and shared between mammals and birds (*5*). In the human larynx, which has been most intensively studied, VF tissue properties, aerodynamic forces and neuromuscular control of laryngeal and VF geometry all interact to ultimately control VF kinematics and glottal flow that set crucial acoustic source parameters such as fundamental frequency, registers, and spectral slope (*1*-*4*, *6*). In contrast to humans, birds are not as well-studied but have the significant advantage that the entire neuromechanical motor control loop of their voice production is experimentally accessible (*7*, *8*). Furthermore the process of song motor sequence learning in songbirds can be monitored individually and shares many parallels with human speech acquisition (*9*–*12*). Thus birds present a tractable model system for *(i)* studying the physics and learned motor control of voiced sound production and *(ii)* to inform on basic principles of MEAD systems control.

Causally linking descending motor control to voiced sound production requires computational biophysical models to explore the high dimensional control space (*13*). High-fidelity continuum models include the full fluid-structure-acoustics interaction (FSAI) complexity of voiced sound production in anatomically realistic geometries of vocal fold and tract (*4*, *14*–*21*). These models are critically needed when realistic representations of voice physiology and biomechanics are essential, such as in the clinical management of voice disorders (*1, 4*), or understanding motor control of human (*22–24*) or avian (*25*) voiced sound production.

However, the continuum model approach has not yet been applied to voiced sound production in avian model systems. Additionally, the approach in general currently faces three key challenges (*1, 4*). *(i)* The implementation itself of complex FSAI models capable of resolving the large range of spatial and time scales remains challenging (*14*-*21*). On the experimental side, although partial physiological datasets have been collected in mammals (*1*, *2, 23*, *24*), we *(ii)* currently lack quantification of all critical physiological parameters in single individuals. This would include parameterization of *(a)* anatomical boundary conditions and (*b*) VF tissue properties, and time-resolved quantification of *(c)* flow, pressure and VF posturing changes due to motor control, and *(d)* VF shape within single oscillations. Obtaining these complete individual-based datasets would lift the last critical roadblock and *(ii)* allow experimental validation of FSAI model predictions to strengthen confidence in the continuum model approach.

Here we resolve the above challenges in an avian model system and implemented and experimentally test predictions of a physics-based, high-fidelity three-dimensional (3D) continuum model of voiced sound production. First, we implemented a 3D immersed-boundary (IBM), finite-element method based fluid-structure interaction solver (See methods), where airflow was governed by 3D, unsteady, viscous, incompressible Navier–Stokes equations and dynamics of the avian VF analog were governed by the Navier equation. The fluid and structure solvers were explicitly coupled through a Lagrangian interface where airway and vibrating structures contacted. To allow rapid simulations when resolving the large range of spatial and time scales in simulating voice production, we used the Message-Passing-Interface parallelization method (*26*), combined with IBM as an advanced numerical method designed especially for simulating moving boundaries. By carrying out simulations on Cartesian grids, IBM circumvents complicated re-meshing algorithms in conventional body-fitted mesh methods and can deal with complex moving boundaries.

To address the second challenge, i.e., quantification of all critical physiological parameters in single individuals, we focused on rock pigeons, where we previously (*5*, *27*) achieved high-quality kinematic data of the avian VF analog, the lateral vibratory masses (LVMs, Fig. 1A). We quantified syringeal anatomy for five individuals using DiceCT scans and implemented high-resolution finite-element meshes of the syrinx (Fig. 1A). To mimic the anatomical constraint of the trachea rings on the motion of the LVMs, zero displacement boundary conditions were applied at mesh nodes where tracheal rings were located. Unilateral LVMs consisted of 1,988-4,620 and 2,159-4,693 nodes and 8,101-18,657 and 9,838-19,131 elements in the solid domain for left and right LVM respectively (Table S1). To parameterize LVM tissue elasticity in each individual, we developed a novel non-invasive, combined experimental/simulation approach. We experimentally induced static LVM displacement by stepwise increase of the transmural pressure (Fig. 1B). We subsequently used the LVM finite-element model combined with genetic-algorithm based optimization (*28*) to simulate LVM displacement as a function of transmural pressure (Fig 1C) and find the elastic modulus (EM) value with the smallest difference between experiment and simulation (Fig 1C). These values were used for further dynamic simulation and ranged from 1.8 to 4.0 kPa for the five individuals (Table S1).

**Figure 1.**
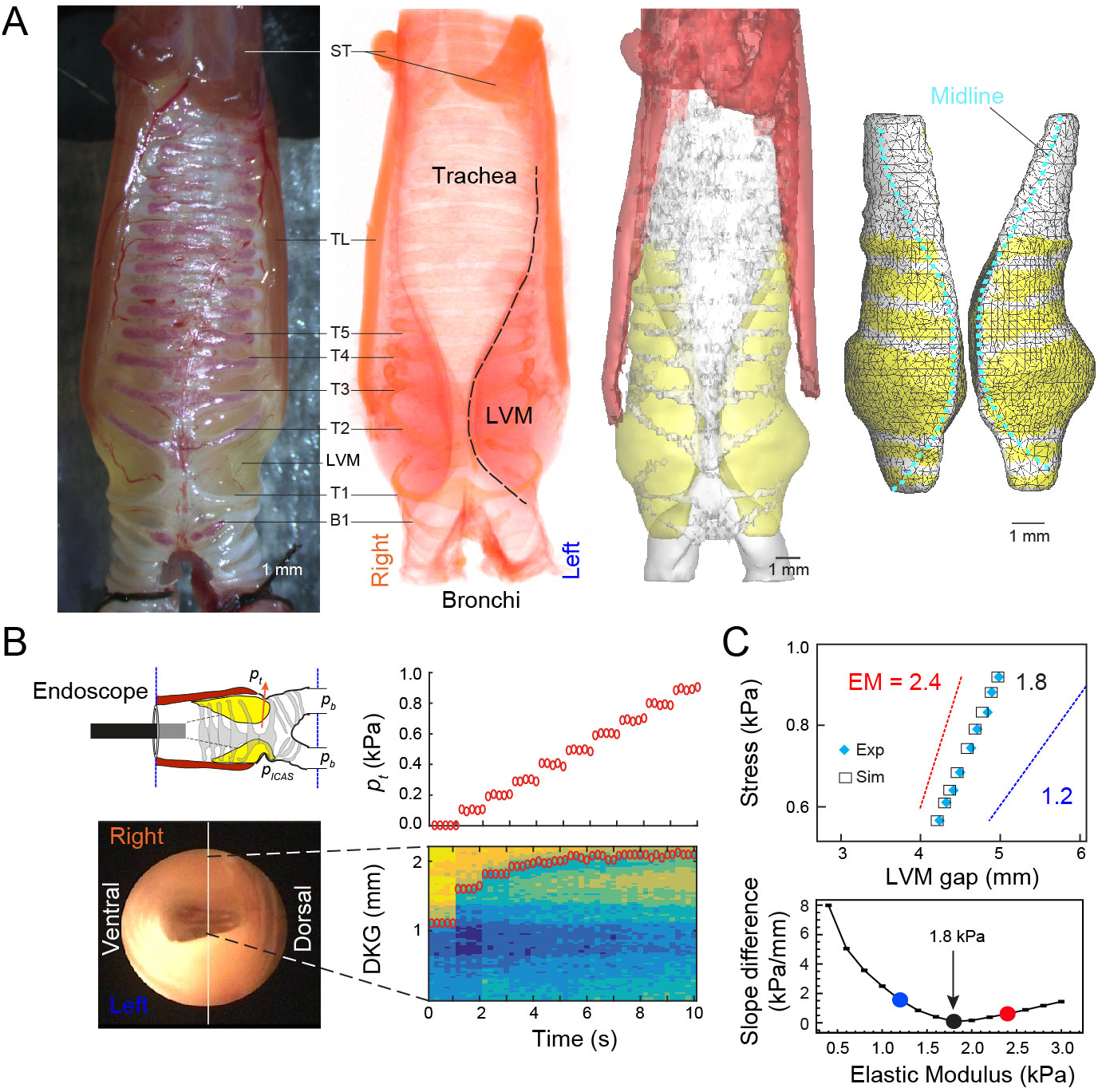
Parameterization of vocal organ geometry and tissue properties. **(A)** Workflow to parameterize FSAI model geometry (Subject P1), with from left to right: 1) photo of syrinx mounted in *in vitro* experimental setup, 2) voltex rendering of iodine contrasted µCT scan showing the bilateral Lateral Vibratory Masses (LVM), 3) 3D anatomy, and 4) Finite Element Mesh of LVM solid domain. Yellow and white mesh elements on LVM outer surface have free and zero displacement boundary conditions, respectively. (B) Tissue properties were determined by a combined experimental and modeling approach. A stepwise increase of (top) transmural pressure *p*_t_ caused a stepwise sideways displacement of the LVMs (bottom) as viewed by digital kymogram (DKG) along the white vertical line in the endoscopic image (left). **(C)** We used a reverse engineering approach to determine the LVM elastic modulus (EM) for each individual. Top: stress-gap width curve of experimental (blue diamonds) and modeled displacement for LVMs with EM=1.8 kPa (black squares). Also indicated are simulated aligned gap widths of EM=1.2 (blue line) and 2.4 kPa (red line). Bottom: Minimal difference between experimental and simulated data indicates occurs at EM = 1.8 kPa for this individual. B1, bronchial ring; T1,…, T5, tracheal rings; LVM, Lateral Vibratory Membrane; *p*_t_, transmural pressure TL, tracheolateralis muscle; ST, sternotrachealis muscle.

To quantify flow, pressure and LVM posturing changes due to motor control, we applied boundary conditions where LVM vibration was obtained reliably previously (*5*, *27*) to both the experimental preparation and FSAI model (*p*_b_ = 1.0 kPa, *p*_icas_ = 0.5 kPa). Both the *in vitro* syrinx and corresponding individual FSAI models demonstrated self-sustained stable oscillations in all five cases (Fig 2; Movie S1). To quantifying the time-resolved VF shape within oscillations, we took advantage of both the unique coronal view offered by the pigeon syrinx (*5*) and the lack of a dorsoventral vibrational component (*27*), to quantify the time-resolved syringeal or glottal opening as a function of caudo-cranial position, i.e. a coronal glottovibrogram (cGVG, Fig 2C).

**Figure 2.**
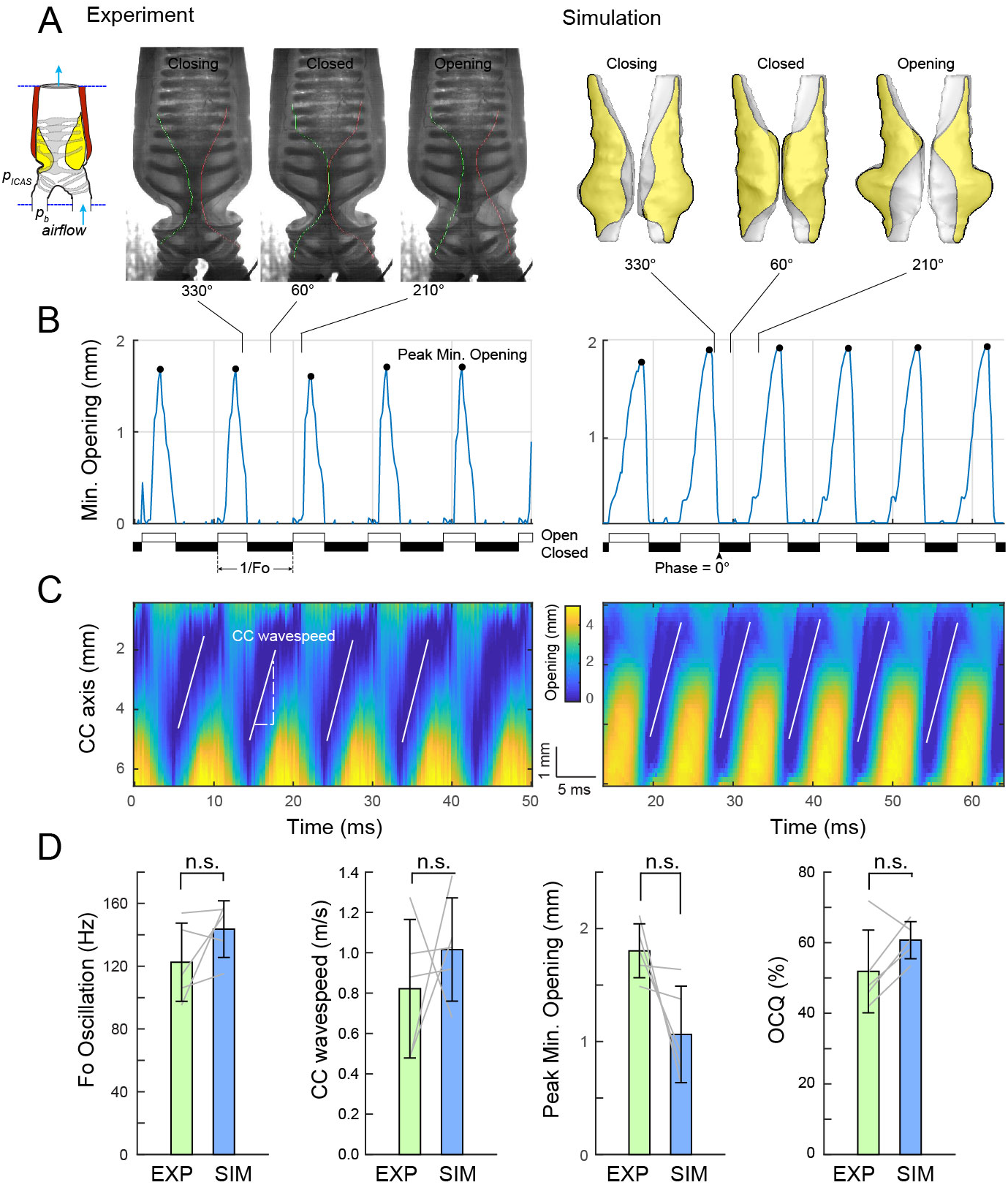
FSAI simulation accurately predicts key features of LVM kinematics. (A) Trans-illuminated syrinx in experiment (EXP, left) and simulation (SIM, right) showing the LVM shape in coronal plane at 330, 60 and 210° of the vibratory cycle, where 0° is defined as the first frame of full closure. (B) The minimal glottal opening shows when the glottis in open and closed (white and black horizontal bars) and vibratory frequency *f*o. The open-closed quotient (OCQ) is the ratio between duration of open and closed glottis. (C) Coronal glottovibrogram (cGVG) showing time-resolved glottal opening along caudocranial axis of 5 oscillations. The white lines indicate the CC wavespeed regression slopes based on the closed glottis. (D) Observed EXP data and SIM predictions are not significantly different for the above indicated four key kinematic parameters.

To rigorously test the predictions of the FSAI model during dynamic simulation (the third challenge), we used a blinded procedure where the modeling team (WJ, QX, XZ) had access only to geometry, static loading test and pressure boundary conditions, but was blind to all other data of the experimental procedures (performed by JHR, CPHE). We compared key vibratory and acoustic predictions by the simulations (SIM) to the behavior in the experiment (EXP). The cGVG allowed for comparing four key parameters describing vibratory kinematics: fundamental frequency of the vibration (*f*_o_), the speed of the caudocranial (CC) wave, peak of the minimal glottal opening, and the open-closed quotient (OCQ). In EXP, *f*_o_ of the LVM vibration measured 123±25 Hz (median: 114 Hz; range: 95-154 Hz, N=5), CC speed was 0.82±0.34 m/s (median: 0.88 m/s; range: 0.48-1.27 m/s, N=5), peak minimal opening was 1.80±0.24 mm (median: 1.80 mm; range: 1.49-2.11 mm; N=5) and OCQ was 1.3±0.7 (median: 1.1, range: 0.7-2.5; N=5). In SIM, *f*_o_ of the LVM vibration measured 143±18 Hz (median: 152 Hz; range: 115-158 Hz, N=5), CC speed was 1.02±0.26 m/s (median: 1.03 m/s; range: 0.68-1.38 m/s, N=5), peak minimal opening was 1.06±0.43 mm (median: 0.90 mm; range: 0.62-1.64 mm; N=5) and OCQ was 1.6±0.3 (median: 1.6, range: 1.1-2.1; N=5). At group level, all predicted values were not significantly different from the experimental values with *p*=0.18, 0.50, 0.06 and 0.11 respectively (Heteroscedastic two-tailed paired t-test, N=5) (Fig 2D). We furthermore compared the time-resolved LVM shape between EXP and SIM at fixed phases within an oscillatory cycle (Fig 3). During the closed phase (0-120°) the simulated LVM shape matched EXP very well and was never significantly different from the bootstrapped shape (p<0.01, see methods). The only observed significant (p>0.05, 2-sample Kolmogorov-Smirnov test) discrepancy occurred during late closed/early opening in three subjects on one side, where the LVM mass tended to move ~0.5 mm more cranial (superior in human anatomical terminology) in EXP compared to SIM (Fig 3BC). Taken together, our FSAI model accurately predicted key kinematic parameters of LVM motion.

**Figure 3.**
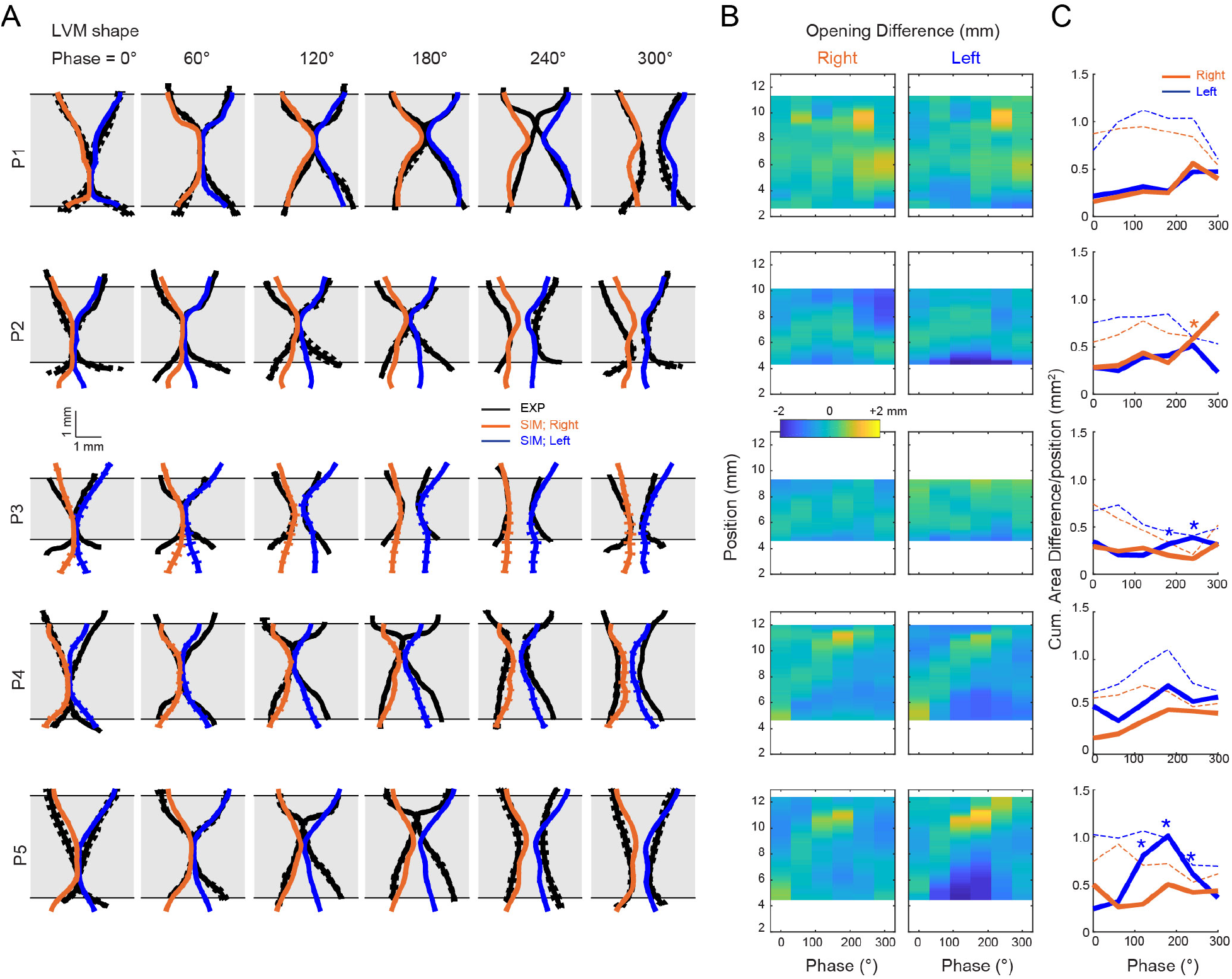
FSAI simulation accurately predicts within-cycle LVM shape. (A) Bilateral LVM shape of EXP and SIM in coronal plane at 0, 60,120, 180, 240, 300 degrees averaged over 5 cycles (mean (solid lines) ± S.D (dotted lines). (B) Lateral position difference between EXP and SIM left and right LVMs and (C) unsigned cumulative area difference between EXP and SIM shows that the predicted shapes are very similar to the observed shapes. The most pronounced discrepancies occur during late closed/early opening. Dotted lines are mean. Asterisks indicate significant difference (p>0.01) between SIM data and 1000x bootstrapped EXP shapes for 2-sample Kolmogorov-Smirnov tests.

Lastly, we compared SIM prediction and EXP observation of two additional key acoustic parameters specifying a sound source – in addition to *f*_o_ (Fig 2D) -, namely source level and spectral slope and these did not differ significantly (Fig 4A). Because the simulated LVM vibratory kinematics matched our observations, we exploited our FSAI models to calculate parameters that could not be quantified in the current experiments, such as spatiotemporal pressure and flow velocity distributions over the vibratory cycle (Fig 4B-E, Movie S2). The convergent LVM shape during opening causes high glottal pressures (~0.9 kPa) that transfer 23.4±15.7 µJ of positive energy (N=5) from flow into LVM (Fig 4F), facilitating opening. When maximum opening is reached at 300° phase, the LVMs are straight (0° angle in Fig 4E) reducing glottal pressure. During early closing the inferior LVM edge is moving inwards causing an energy transfer of 7.2±2.2 µJ back into the flow. Consecutively, the LVMs close the glottis by moving together in a divergent shape causing rapid pressure reduction driven by elastic forces (Arrow in Fig 4E). Furthermore, flow inertia in the trachea causes negative pressures (−0.52±0.23 kPa) near the glottis exit prior to full closure (blue region in Fig 4D), facilitating closing. Interestingly, another positive energy transfer to LVMs is observed nearly the end of the cycle. Our avian data thus show that two primary factors contribute to the pressure asymmetry during VF opening and closing that drive self-sustained oscillation: *i)* an alternating convergent/divergent medial surface profile and *ii)* airflow inertia, corroborating earlier model predictions (*29*) and measurements on (hemi)larynges (*30*) in mammalian voice production.

**Figure 4.**
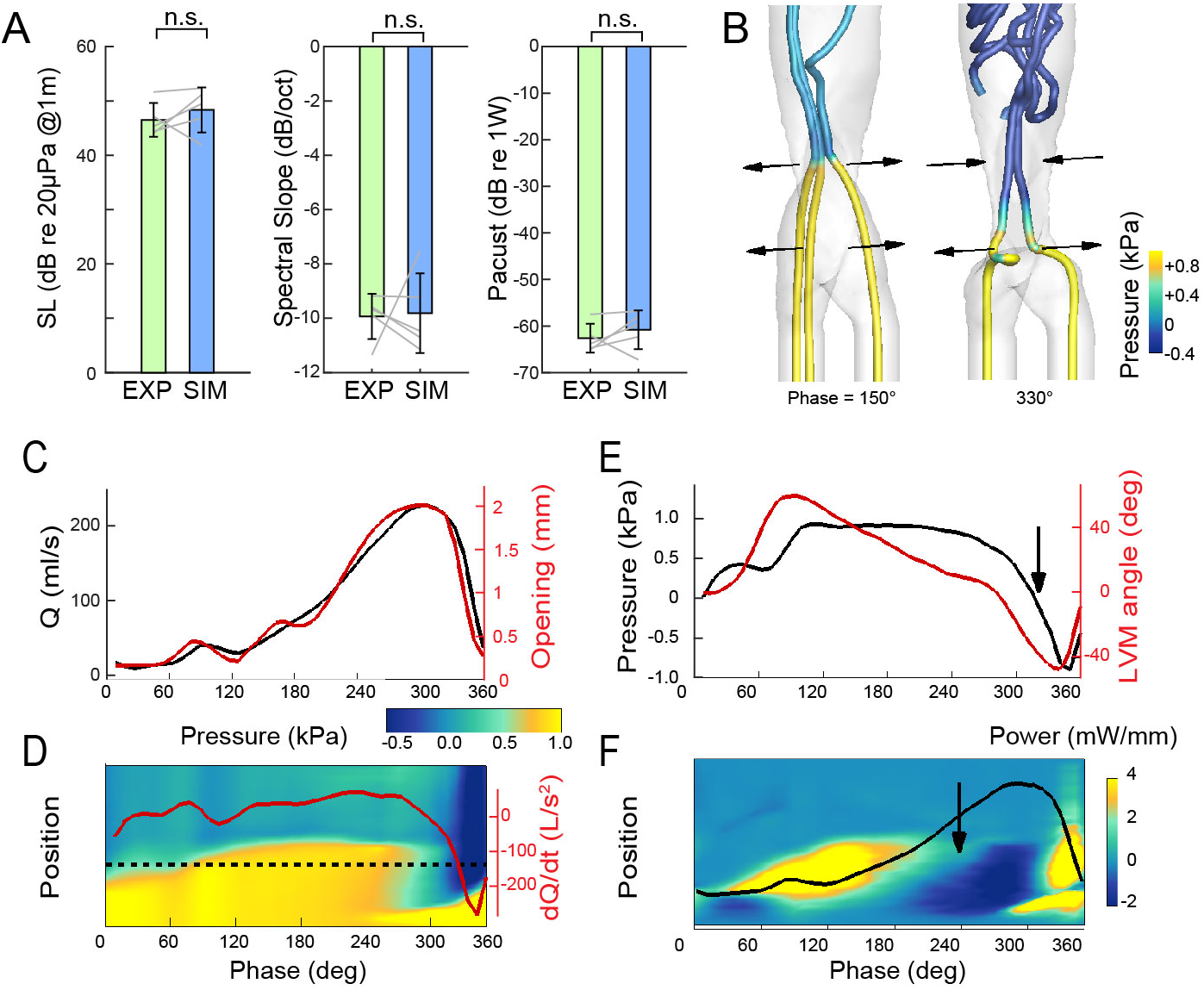
FSAI predictions of acoustics and spatiotemporal pressure and power profiles. (A) The SIM accurately predicted key acoustic parameters; source level (SL), spectral slope and acoustic power. (B) 3D flow inside the syrinx as indicated by flow streamlines (pressure color-contoured). Horizontal arrows indicate the motion direction of the LVMs. (C) Flow rate Q (black line) correlates strongly with glottal opening (red). (D) Spatiotemporal pressure distribution along the airway centerline over a cycle with superimposed dQ/dt (red solid line) as a proxy for sound pressure. (E) Glottal pressure (black line) evaluated at the horizontal dotted line in (D) with LVM angle (red line). (F) Spatiotemporal power transfer distribution from flow to LVM evaluated along the airway centerline with superimposed flow Q (black solid line).

Our FSAI model accurately predicted key performance traits of tissue motion and acoustics driven solely by physiological parameters (static geometry, tissue elasticity and boundary conditions) and without optimization of either geometry or material properties parameterization on dynamic performance (*20*). While the FSAI model implementation itself is complex (*14*–*21*), the inputs are simple and have, most importantly, directly measurable physiological, material and geometrical properties. Measurements of VF material properties (*31*, *32*), initial configuration, initial stress and detailed flow-induced 3D VF motion (*33*) have been achieved separately in human and mammalian model systems, but complete physiological data sets have not been obtained in these clades, nor in any birds, in single individuals nor consecutively used to thoroughly test FASI model predictions in a blinded approach. Recent studies encouragingly suggested that realistic continuum 3D models lead to more robust VF dynamics compared to 1D and 2D models (*34*). Our data shows that they also lead to accurate predictions performance. Our results therefore strongly support the continuum model approach as a critical step in integrating *in vitro, ex vivo and in vivo* experimental data with brute force computational approaches towards a causal model of motor control of voiced sound production applicable to birds and mammals.

Continuum models currently allow only limited systematic parameters exploration because they are computationally expensive (*21*). In contrast, phenomenological reduced order (RO) models - that simplify VFs to one or two masses (*35*, *36*)-allow broad exploration of a simplified control space with less computational power and as such have been used widely to capture the dynamics of mammalian (*37*) and avian voiced sound production (*13*, *38*–*40*). However, because RO models are not first-principle physics models they essentially lack physiological representation of tissue properties and geometry (*1*, *4*, *37*). Reducing computational costs and time is thus crucial for making continuum models useful in experimental or clinical settings (*1*). Rapid developments in machine learning tools combined with supercomputing provide promising avenues to predict complex flow solvers (*41*, *42*) that could significantly reduce computational power and allow elaborate parameter exploration.

All vocal behaviors result from system-wide interactions between the nervous system, body, and surrounding environment (*43, 44*). Understanding the activity of neural circuitry controlling voice can therefore only be understood by embodied models that include realistic biomechanics of sound generation and musculoskeletal actuation of the vocal organ (*8*, *45, 46*). During postnatal vocal development, gradual body changes, such as lung growth or VF stiffness in marmosets (*46, 47*) or increased muscle speed in songbirds (*48*), can drive changes in vocal behavior. Embodied human VF models have direct clinical relevance to pathological physical behaviors with abnormal vocal output (*1*) and patient specific model-assisted phonosurgery (*49*, *50*), because most laryngeal human voice disorders are caused by changes in VF geometry, structural integrity, or kinematics (*4*). The embodied approach to voice production as presented here thus improves our understanding how neural mechanisms and biomechanics interact to drive vocal behavior in vertebrates.

## Acknowledgements

The authors with to thank T Christensen, S Jakobsen, P Martensen and F Mortensen for technical support, I Adam for discussion, and C Herbst and J Rattcliffe for comments on the manuscript.

## Author contributions

WJ, JHR, QX, XZ, CPHE designed research; WJ, JHR, MD, QX, XZ, CPHE performed research, with the modeling team consisting of WJ, QX and XZ, and experimental team of JHR and CPHE; CT scans were performed by JHR and MD; QX, XZ, CE contributed new reagents or analytic tools. WJ, JHR, QX, XZ, CPHE analyzed data; CPHE wrote the first draft; All authors read and approved the final manuscript.

## Data and materials availability

All data and code used for analysis are available upon request.

