## Supplemental Material for "High-fidelity continuum modeling predicts avian voiced sound production"

**List of Supplementary Materials:**

Materials and Methods

Table S1

Figure S1

Movie S1-2

### MATERIALS AND METHODS

#### Subjects

We used 6 adult domestic pigeons (*Columba livia*) obtained from local breeders. The subjects were kept in an outdoor aviary (3x6x2 meters) at the University of Southern Denmark with access to food and water *ad libitum*. All procedures were carried out in accordance with the Danish Animal Experiments Inspectorate (Copenhagen, Denmark).

#### Experimental Setup

Subjects were euthanized by overdosing with Isoflurane (Baxter Medicals, Il, USA) and the syrinx was extracted from the primary bronchi up to including ~10 cm of trachea. The syrinx was mounted to tracheal and bronchial connectors in the experimental setup described in detail in (*5*, *26*). In brief, this setup allows fine-control of bronchial ($p_b$) and air sac ($p_{as}$) pressure, flow, temperature, and simultaneous high-speed imaging from sagittal and endoscopic tracheal views. Both pressures were controlled with a dual valve differential pressure PID controller (0–10 kPa, model PCD, Alicat Scientific, AZ, USA), referenced to atmospheric pressure. Bronchial mass flow was measured with flow sensors (0-3 liters/min, PMF 2104, Posifa Microsystems, San Jose, USA). Sound was recorded with a ½-inch pressure microphone (model 46AD with preamplifier type 26AH, G.R.A.S., Denmark), amplified and high-pass filtered (10 Hz, 3-pole Butterworth filter, model 12AQ, G.R.A.S.). The microphone sensitivity was measured before each experiment (sound calibrator model 42AB, G.R.A.S.). The microphone was placed 5-10 cm from the tracheal connector outlet on a 45° angle to avoid air jets from the tracheal outlet. Microphone, pressure and flow signals were low-pass filtered (10 kHz, EF120, Thor labs, NJ, USA) and digitized at 48 kHz (USB 6259, 16 bit, National Instruments, Austin, TX, USA) together with TTL trigger to synchronization for the camera systems. All control and analysis software was written in Matlab (Mathworks).

**Experimental Procedures**

*1. LVM tissue properties*: After mounting we first measured the stress-stain relationship of the LVM tissue using a static loading test. We imaged the static deformation of the LVMs using a 1.2 mm flexible endoscope (Scholly, Germany) attached to a videokymographic (VKG) system (Videokymographic camera 2156, Cymo B.V., The Netherlands), which combines a high-speed linescan camera (7,200 lines/sec) with a full frame CMOS camera (25 frames/sec), while stepwise increasing transmural pressure every 1.5 s from 0-2 kPa (in increments of 0.1 kPa) corresponding to the *in vivo* pressure range (Beckers, 2003; Gaunt, Gaunt, & Casey, 1982; Elemans 2008). To prevent airflow through the syrinx, the trachea was closed. The analog video output from the VKG was digitized together with the microphone and a synchronization signal using a video capturing device (Intensity Extreme, Black Magic Design, Australia). We constructed digital kymograms perpendicular across the LVM. The edge position was quantified using outwards-in threshold detection of the edge. The length of dorsoventral glottis at mid LVM position was determined from each CT scan and allowed us to calibrate LVM displacement from the DKG in mm. For statistical purpose, five 5 technical replicates were made for each transmural pressure setting.

*2. LVM kinematics during voiced sound production*. Second, we quantified the time-resolved motion of the LVMs, flow, pressure and sound during phonation. We applied the boundary conditions of $p_b$ = 1.0 kPa, $p_{as}$ = 0.5 kPa, where LVM oscillations reliably occurred (*5*, *26*). The resulting positive transmural pressure ($p_t = p_b - p_{as}$) mimics activation of the TL muscle *(40)*. A powerful, stable light source (1700 Lumen white LED powered by PS23023, HQ Power amplifier, Belgium) trans-illuminated the syrinx from dorsal allowed us to capture LVM motion with a high-speed camera (MotionPro-X4, 12 bit CMOS sensor, Integrated Design Tools, Inc.; 4,000 frames/sec) mounted on a stereomicroscope (M165-FC, Leica Microsystems). Before and after experiments we made a photo of the mounted preparation with a Leica DFC400 digital camera mounted on the M165-FC stereoscope. LVM shape was hand-traced in all specimens on 150-200 selected consecutive high-speed images (covering at least 5 full vibratory cycles) using a custom-made Matlab graphical user interface.

3. *Syrinx geometry based on DiceCT scan*. Lastly, after experiments were completed we mounted the syrinx with Austerlitz insect pins (Fine Science Tools, Germany) onto a 5 mm thick sheet of stiff silicon, taking care to mimic the exact geometry of the syrinx during the experiments based on earlier photos. The preparation was submerged into phosphate buffered saline solution (PBS) with 4% paraformaldehyde (PFA) for at least 24 hours to fix the geometry of each preparation. After fixation, the preparations were transferred to PBS and kept at 5C. After mounting, after fixation and prior to scanning we made photos with a Leica DFC400 digital camera mounted on the M165-FC stereoscope to quantify shrinkage and shape changes. One preparation was omitted for further analysis as it twisted after fixation compared to the experimental situation. Preparations were removed from the silicon base and stained in 10% IKI in demineralized water and rotated at 5 RPM for 24 hours twice (*51, 52*). Preparations were washed in demineralized water 8 times just prior to being transferred into a microCT scanner (vivaCT 40, Scanco Medical A., Brüttisellen, Switzerland). Scanning settings were 70 kV and 57

μA. After scanning, the 3D reconstruction cubic voxel size measured 10.5*10.5*10.5 μm3 (2048*2048*2048 pixels) with 32-bit gray-levels.

**Blinding Procedure and Data Disclosure**

To eliminate the chance of fitting model predictions on experimental data, we implemented the following blinding procedure. The experimental team (JHR, CPHE) performed the experimental procedures listed above. Afterwards, the modeling team (WJ, QX, XZ) was given access only to the DiceCT scan, static loading test (LVM displacement and transmural pressure) and asked to predict key performance traits of LVM kinematics (glottovibrogram) and acoustic waveforms during voiced sound production under listed posturing and pressure boundary conditions. After the simulations of all preparations were completed all experimental data was disclosed and analyzed.

**Fluid-Structure Acoustic Interaction (FSAI) Model**

The computational solver was built upon an immersed-boundary-finite-element method based fluid-structure interaction solver (*14-16*). The airflow was governed by the three-dimensional, unsteady, viscous, incompressible Navier–Stokes equations:

$$\nabla \cdot \vec{U} = 0$$

$$\frac{\partial \vec{U}}{\partial t} + \left(\vec{U} \cdot \nabla\right)\vec{U} = -\frac{1}{\rho_0}\nabla \mathrm{P} + \upsilon_0 \nabla^2 \vec{U} \qquad (1)$$

where $\vec{U}, \rho_0, P, \upsilon_0$ are the incompressible flow velocity, density, pressure and kinematic viscosity, respectively. The LVM dynamics was governed by the Navier equation with a linear stress-strain relationship:

$$\rho_{tiss}\frac{\partial^2\vec{d}}{\partial t^2} = \bar{\bar{\sigma}}\cdot\overleftarrow{\nabla} + \rho_{tiss}\vec{f} \tag{2}$$

where $\rho_{tiss}$ is the tissue density, $\vec{d}$ is the displacement, $\bar{\bar{\sigma}}$ is the stress tensor, and $\vec{f}$ is the body force. Details regarding the numerical algorithm of the flow and solid solvers can be found in (*16*).

The fluid and structure solvers were explicitly coupled through a Lagrangian interface where airway and LVMs contact. In each iteration, the fluid solver was first marched by one step with the existing deformed shape and velocities of the LVM surface as the boundary conditions. The forces at the LVM surface were then calculated with the new fluid pressure. At the second step, the solid solver was marched by one step with the updated surface traction. The deformation and velocities of the LVM surface formed the new fluid boundary condition, so that the fluid solver and solid solver were coupled. The acoustic pressure was first calculated by the linearized perturbed compressible equation (LPCE) based on the hydrodynamic/acoustic splitting method (*53, 54*):

$$\begin{gathered}
\frac{\partial\rho'}{\partial t} + \left(\vec{U}\cdot\nabla\right)\rho' + \rho_0(\nabla\cdot\vec{u}') = 0 \\
\frac{\partial\vec{u}'}{\partial t} + \nabla\left(\vec{u}'\cdot\vec{U}\right) + \frac{1}{\rho_0}\nabla p' = 0 \\
\frac{\partial p'}{\partial t} + \left(\vec{U}\cdot\nabla\right)p' + \gamma P(\nabla\cdot\vec{u}') + (\vec{u}'\cdot\nabla)P = -\frac{DP}{Dt}
\end{gathered} \tag{3}$$

where $\rho'$, $\vec{u}'$, $p'$ are compressible perturbed flow density, velocity, pressure, $\gamma$ is the ratio of the specific heats.

With this splitting method, the total velocity/pressure of the flow would be the sum of the incompressible flow velocity/pressure and acoustic velocity/pressure perturbation; hence the acoustic pressure obtained from incompressible fluid field also forms a loading on LVM surface mesh. However, the calculation of the LPCE showed that the root-mean-value of the acoustic flow rate was only 0.9% of the incompressible flow rate, suggesting a weak coupling effect between the acoustic field and incompressible flow field. Therefore, the linear source-filter theory (*2*) was applied which assumes that the sound radiation in the tract is linearly coupled with the source generation in the syrinx so that the acoustic pressure does not affect vibrations. Because the dominant sound source of voice is a monopole source, the far-field acoustic pressure was calculated as below:

$$p' = \frac{\rho}{4\pi r}\frac{dQ}{dt} \tag{4}$$

where $p'$ is the acoustic pressure, $\rho$ is the air density, $r$ is the distance from sound source to the microphone and $dQ/dt$ stands for the time derivative of flow rate.

Power transfer from airflow to the vibration masses over a vibration cycle was calculated by multiplying the flow pressure force with the normal components of velocity vectors.

**Parameterization of Vocal organ Geometry**

The geometry of the airway and LVMs was reconstructed manually from the DiceCT scan of each specimen in Mimics (Materialise, Leuven, Belgium). We used every tenth slice of the original scan in this segmentation process. The surface mesh of the airway and LVMs was exported from Mimics. The LVM tetrahedral mesh for finite-element analysis was generated using ANSYS or Gmsh and contained around 14,000 tetrahedral elements for each LVMs. Based on the anatomy, several places on the LVM surface were connected to syringeal solid structures, such as ossified tracheal rings, and therefore the motion at these places was constrained in the simulation. All other nodes were free to move and a traction boundary condition was applied.

**Simulation Implementation and Conditions**

The vocal organ mesh was immersed in a computational domain that was about 12×10×40 mm in dimensions and straight tubes were added to each bronchial inlet and tracheal outlet to avoid reverse flows. The domain was discretized with a non-uniform 64×48×128 Cartesian mesh with the highest grid density around the LVMs. A non-penetration non-slip boundary condition was applied at the airway wall. The part of the airway wall contacting the LVMs was the interface of the fluid-structure interactions on which the deformation and velocity of the LVM surface were transferred to the fluid solver and the fluid pressure on the LVM surface was transferred to the solid solver. The air density was set to 1.1455 kg/m$^3$ at 37°C. We used a kinematic viscosity value of 6.6×10$^{-5}$ m$^2$/s (at 37°C) - four times the normal value - to reduce high frequency turbulence in the simulations, which significantly reduces the computational costs (*34*).

We assumed the LVM in pigeons consists of isotropic material based on earlier histological sections (*55*). We assumed incompressible material properties and the Poisson ratio ($v$) was set to be 0.46 to avoid the singularity problem at $v$=0.5. The shear modulus ($G$) was obtained from the relationship of $G=E/2/(1+v)$ for isotropic materials. Tissue density was assumed to be 900 kg/m$^3$, equal to the density of fat.

For the dynamic simulations a time step of about 1.0 μs was utilized in the fluid and solid solvers. Two artificial non-slip and non-penetrable collision planes were 80 μm off the medial plane to enforce a finite 160 μm minimum gap between the two LVMs during closure. The LVM position was not allowed to exceed the corresponding collision plane during collision, to prevent failure of the solver due to the non-conserved mass in each zone. For each individual, the simulation was performed on 16 processors and took about 1.5 days per vibration cycle. We obtained at least five cycles per individual.

### Parameterization of LVM Elastic Moduli

To parameterize tissue elasticity for each individual, we developed a novel combined experimental/simulation approach. We used the finite-element LVM model to simulate LVM displacement as a function of transmural pressure as obtained in the experiment listed above for a range of elastic moduli values. Uniform bronchial pressure steps were applied on the inner LVM surface and the maximum LVM gap was obtained for each pressure step. For each preparation, different elastic modulus values generated by a genetic algorithm-based optimization method (*27*) were randomly assigned to each static analysis solver, which resulted in a different displacement-transmural pressure slope. We determined the elastic modulus of the LVMs for each preparation in an iterative process by minimizing the difference between the slopes of LVM displacement versus $p_t$ measured in the experiment with the 3D mesh model.

### EXP and SIM Data Analysis

*Kinematic Analysis:* To compare LVM kinematics we calculated the time-resolved glottal opening (i.e. a glottovibrogram) from the left and right LVM position as measured in the highspeed video (EXP) and along the midline of the simulations (SIM). In this spatiotemporal matrix of glottal opening, we calculated the time-resolved minimum opening to detected values of complete closure, i.e., glottal opening = 0 μm for EXP and <100 μm for SIM over five cycles. The first instance of closure (defined as $\varphi$=0°) was used to calculate instantaneous vibration period and fundamental frequency per cycle. The Open-Closed Quotient (OCQ) was defined as the ratio of time open over time closed per cycle. The caudo-cranial (CC) wavespeed was calculated by linear regression of all data points of complete closure as a function of CC position and time in a cycle. For all kinematics parameters the mean over 5 cycles was used for further comparison.

To compare the LVM shape between EXP and SIM in more detail we first aligned EXP and SIM data over the midline and focused on the range where data was present for both EXP and SIM along the entire vibratory cycle (grey boxes in Fig 3). We calculated the LVM opening difference along the CC axis position between the mean shape of EXP and SIM over 5 consecutive cycles at phase $\varphi$=0, 60, 120, 180, 240 and 300°. To statistically compare LVM shapes we first calculated the cumulative difference between EXP and SIM opening along the CC axes (i.e. the area between LVM shapes). We then repeated this process 1000 times against a randomized EXP shape to create a distribution of mean area difference and tested whether the LVM shape and random shape were drawn from same continuous distribution using a two-sample Kolmogorov-Smirnov test per phase step.

*Acoustical Analysis*: To compare the acoustics of the model with the experimental obtained values we first resampled the SIM signal from about 1.2 MHz to 48 kHz using the resample function in Matlab. We low-pass filtered EXP and SIM sound signal at 20 kHz with a 3$^{th}$ order butterworth filter with zero-phase shift implementation (filtfilt.m function). Source level was defined at 1 meter from the source as $SL = 20\log(p/p_{ref}) + TL$, with $p$ the RMS pressure and $p_{ref}$ =20 µPa, transmission loss $TL = 20*\log10(d)$ with distance $d$=12 cm. Spectral slope was calculated following (*6*). We first detected all peaks in a spectrum based of all 5 cycles. Spectral slope was defined as the slope of the linear regression of amplitude over log10 frequency of all detected peaks multiplied by log10(2).

*Statistical Analysis:* One preparation was omitted for further analysis as fixation caused it to twist compared to the experimental situation. All data is presented as mean ± S.D. and comparisons were done with paired-sample t-tests at significance level $p$=0.05, unless mentioned otherwise.

**Table S1. LVM finite element model statistics and elastic moduli per individual.**

| | Nodes | | Elements | | EM |
|---|---|---|---|---|---|
| ID | Left LVM | Right LVM | Left LVM | Right LVM | kPa |
| P1 | 3988 | 3835 | 15737 | 15190 | 1.69 |
| P2 | 1988 | 2278 | 8101 | 9838 | 1.8 |
| P3 | 2686 | 2159 | 11402 | 8532 | 4 |
| P4 | 4253 | 4551 | 17062 | 18436 | 2.24 |
| P5 | 4620 | 4693 | 18657 | 19131 | 2.4 |

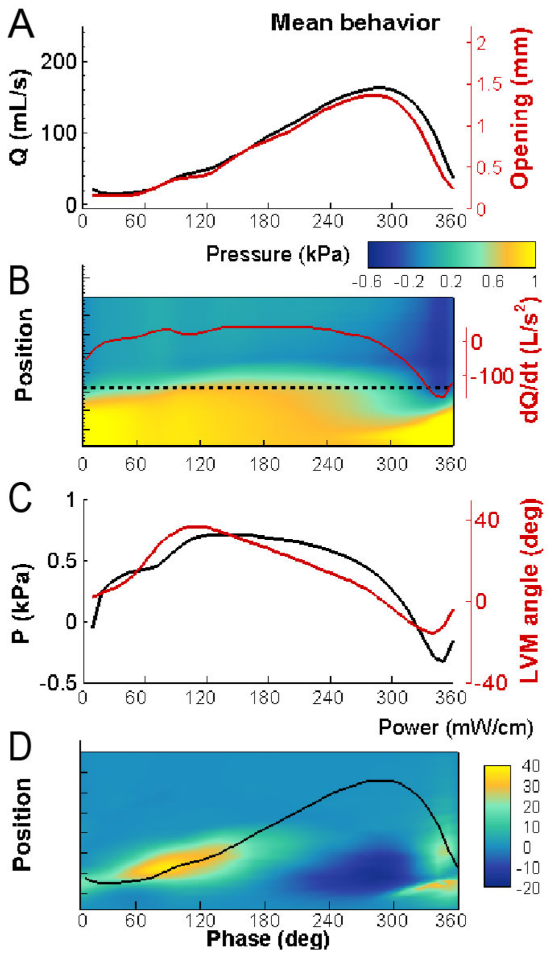


**Figure S1. Mean FSAI prediction of spatiotemporal pressure, power profiles and acoustics.** Mean performance for all 5 individuals. (A) Flow rate Q (black line) correlates strongly with glottal opening (red). (B) Spatiotemporal pressure distribution along the airway centerline over a cycle with superimposed dQ/dt (red solid line) as a proxy for sound pressure. (C) Glottal pressure (black line) evaluated at the horizontal dotted line in (B) with LVM angle (red line). (D) Spatiotemporal power transfer distribution from flow to LVM evaluated along the airway centerline with superimposed flow Q (black solid line).

**Movies**

**Movie S1.** *In vitro* oscillation with corresponding *in silico* oscillations.

**Movie S2.** LVM oscillation of subject P1: (a) fluid region with the streamline contoured by pressure; (b) tissue kinematics contoured by pressure loading (c) power and energy transfer within the two cycles.
